# Red Blood Cell Transfusion is a Non-Canonical Immune Stimulus Characterized by the Suboptimal Induction of CD4^+^ T Cell Help

**DOI:** 10.1101/2025.06.26.661795

**Authors:** Jelena Medved, Abhinav Arneja, Neha Shah, Benjamin N. Hester, Emily D. Burnett, Alexis R. Boscia, Tamara C. Moscovich, Whitney R. Swain, Aditi S. Kodali, Arwen Chandler, Gunnhildur A. Thorkelsdottir, Catherine J. W. Schwarzschild, Rachel J. Muppidi, Iman Cherkaoui, Jessica M. Strand, Michael Trubetskoy, Elizabeth A. Stern-Green, Soraya N. Tarrah, Andria Li, Conrad S. Niebuhr, Juan E. Salazar, Manjula Santhanakrishnan, James C. Zimring, James D. Gorham, Krystalyn E. Hudson, Jeanne E. Hendrickson, Chance John Luckey

## Abstract

Red blood cell (RBC) alloimmunization to non-ABO antigens is a major clinical complication for chronically transfused patients. When exposed to transfused RBCs carrying foreign antigens, some patients generate IgG antibodies that target these antigens, creating potential barriers to future transfusions. Interestingly, other patients produce only IgM antibodies against the same non-ABO antigens, which generally have fewer clinical consequences. Despite the stark differences in their impact, the factors regulating IgM versus IgG production in response to transfused RBCs remain poorly understood. This study explores the balance between IgM and IgG production following transfusion, comparing it to the well-characterized antibody response induced by vaccination in mouse models. By directly assessing antibody levels following RBC transfusion versus Alum-adjuvanted vaccination, we demonstrate that transfusion of RBCs expressing a model antigen is a relatively weak inducer of IgG class switching. Additionally, loss-of-function experiments using CD40L blockade and CD4 depletion confirmed that T cell help is essential for class switching after transfusion but has no effect on IgM production. Most notably, providing supra-physiological levels of T cell help enhanced class switching in a dose-dependent manner after transfusion, whereas vaccination-induced class switching remained unaffected. These findings support a model in which the limited IgG class switching following transfusion stems from suboptimal T cell help compared to vaccination. Furthermore, they suggest that transfusion activates T cells through a non-canonical pathway, distinct from the mechanisms driving immune responses to standard Alum vaccination.

## Introduction

Transfused red blood cells (RBCs) often carry foreign (allogeneic) antigens, and some recipients develop antibodies against these antigens, a process known as RBC alloimmunization.(1) While IgG alloantibodies can lead to severe complications in chronically transfused patients, including life-threatening reactions upon re-exposure to the antigen, others generate only IgM alloantibodies, which are generally not clinically problematic. Despite these stark differences in clinical impact, the mechanisms that determine whether a patient predominantly produces IgM or IgG in response to transfused RBCs remain poorly understood.

In mouse models of RBC alloimmunization, most RBC alloantigens elicit detectable IgM responses but rarely induce IgG class switching.(2) One exception is the HOD mouse model, where donor RBCs express a triple fusion protein that consists of hen egg lysozyme (HEL), ovalbumin (OVA), and human Duffy (HOD).(3) In our previous work, we demonstrated that transfusion of wild-type (WT) mice with HOD RBCs results in the production of both IgM and IgG.(4) However, IgG responses after transfusion were short-lived and of lower-affinity compared to those induced by canonical immune stimuli, such as vaccination. This fast evanescence and reduced affinity of antibodies generated in response to transfusion likely results from insufficient germinal center (GC) formation, which are a hallmark of robust IgG responses observed after vaccination.(4–8) Why transfused RBCs fail to drive robust GCs, high-affinity IgGs, and long-lived IgG responses remains an open question.

Effective adaptive immune responses to canonical T cell-dependent antigens (TD), such as protein-adjuvant vaccination, rely on the presence and function of CD4^+^ T cells.(9–14) Specifically, class-switching to IgG, development of robust GCs, and long-lived high-affinity IgG production all depend critically on provision of CD4^+^ T cell help directly to B cells. CD4^+^ T cell help to B cells is mediated through coordination of multiple pathways, which include direct cell-cell contact interactions and directed secretion of cytokines and other soluble effector molecules.(11,15–20) One of the primary mechanisms via which CD4^+^ T cells provide help to B cells is through cell-cell contact dependent CD40-CD40L interactions.(20–26) CD40, primarily expressed on B cells and antigen-presenting cells, interacts with CD40L, a TNF family member predominantly found on activated CD4^+^ T cells.(21,26,27) This interaction is crucial for class switching, affinity maturation, and sustained antibody responses.(28) Although B cells can be activated without CD40 signaling, key functions such as class switching and GC formation are impaired in its absence.(21,26)

Previous work with mouse models has demonstrated that CD4^+^ T cell help is essential for IgG production in response to transfused RBCs. Depleting CD4^+^ T cells or blocking CD40-CD40L interactions prevents class switching to IgG in response to transfused RBCs, reinforcing the role of T cell-dependent immunity in RBC alloimmunization.(29,30) Although transfusion-driven IgG production critically depends on CD4^+^ T cells, it remains unclear why other CD4^+^ T cell-dependent responses, such as robust GC formation and high-affinity IgG production, are relatively diminished compared to those induced by protein-adjuvant vaccination.

In this study, we expanded on our previous work and investigated efficiency of class switching to IgG and the role of CD4^+^ T cell help following RBC transfusion using the HOD mouse model. To better understand the nature of immune responses against transfused RBCs, we compared and contrasted transfusion-induced antibody responses to those elicited by vaccination with HEL-OVA emulsified with aluminum hydroxide (Alum), which is a well-characterized TD immunization model. By directly comparing the level of antibodies following immunization, we demonstrated that transfusion induces robust IgM production but limited IgG class switching compared to vaccination. Investigation and comparison of CD4^+^ T cell help in transfusion and vaccination driven immune responses revealed further differences between the two immune stimuli. Loss-of-function experiments demonstrated that despite transfusion driven IgG responses requiring CD4^+^ T cells, IgM production is CD4^+^ T cell independent. Through gain-of-function experiments, we showed that transfusion induces CD4^+^ T cell help at suboptimal levels compared to vaccination. Despite sharing features with canonical TD antigens, we observed differences in the role and quality of CD4^+^ T cell help in transfusion driven immune responses when compared directly to vaccination. Taken together, our results highlight the non-canonical nature of transfused RBCs as an immune stimulus.

## Methods

### Mice

C57BL/6J WT, OT-II and CD45.1 mice were purchased from Jackson Laboratories (strain ID 000664, 004194, and 002014 respectively). OT-II mice were crossed with CD45.1 mice. HOD transgenic mice express a triple fusion construct of Hen Egg Lysozyme, Ovalbumin, and the human Duffy RBC antigen selectively on the surface of RBCs, and were generated on the FVB background as described previously.(3) All mice were bred and maintained at the animal facilities of the University of Virginia and Bloodworks Northwest. Mice were immunized between the ages of 8-10 weeks. Experimental groups were age and gender matched. All procedures were approved by Institutional Animal Care and Use Committees at the University of Virginia and at Yale University.

### Murine blood collection

Collection, processing, and storage of HOD blood was performed as described previously.(4,31) Briefly, blood from HOD mice was collected aseptically directly into the anticoagulant citrate phosphate dextrose adenine solution (CPDA-1, Boston Bioproducts IBB-420) by cardiac puncture. The final CPDA-1 concentration was kept at 20% (v/v). Collected HOD blood was leukoreduced using a whole blood cell leukoreduction filter (Pall, AP-4851). Leukoreduced blood was centrifuged at 1200 X *g*, adjusted to a final hematocrit level of 75% and stored at 4°C for 12 days before transfusion.

### HEL-OVA conjugation

HEL-OVA conjugation was done as previously described.(32) Briefly, Ovalbumin (450 mg; Sigma A5503) and HEL (126 mg; Sigma L4919) were dissolved in 18 ml of phosphate buffer (pH 7.5). The solution was centrifuged at 450 X g for 5 min and 14.4 ml of 0.3% glutaraldehyde solution (78.6 μl of glutaraldehyde (Electron Microscopy Sciences 16120) in 24 ml phosphate buffer) was added to the supernatant and mixed for 1 h at room temperature before another centrifugation at 450 X g for 5 min. The supernatant was then dialysed (Thermo 87737) overnight against PBS at 4°C before sterile filtration through 0.2 μm filters (Thermo 564-0020). Protein content was determined using a Micro BCA Protein Assay Kit (Thermo 23235) and aliquots of 1 mg/ml were stored at −20°C until required.

### Immunization and phlebotomy of mice

HOD blood was taken out of storage, pipetted up and down gently to make a uniform suspension of RBCs, and transfused into recipient mice via retro-orbital injection. Each recipient mouse received 100 μl (volume adjusted equivalent of 1 unit of packed RBCs in humans) of stored HOD blood. For vaccination experiments, 100 μg of HEL-OVA conjugate was thoroughly mixed with 100 μl aluminum hydroxide (ALHYDROGEL^→^, vac-alu-250, InvivoGen) and given to recipient mice via IP injections. Blood samples for antibody measurements from immunized mice were collected by submandibular vein puncture and allowed to clot. Samples were then spun at 5,000 rpm for 5 min and the supernatant was transferred to new tubes. This process was then repeated for the second time in order to get sera with no RBC contamination. Sera were stored at −20°C until analysis for anti-RBC alloantibodies.

### Antibody treatment

For CD40L blocking experiments, mice were given 250 μg of the anti-mouse CD40L blocking antibody or isotype control (BioXCell, BE0017-1 or BE0091) 2h-4h before immunization, and subsequently on days 3, 7, and 10 after immunization. For CD40 stimulation experiments, mice received 100 μg of the CD40 activating antibody or isotype control (BioXCell, BE0016-2 or BE0089) 2h-4h before immunization. For CD4 depletion experiments, mice were injected with two doses of 200 μg anti-mouse CD4 antibody or an isotype control (BioXCell,BE0003-1 or BE0090) 4 and 2 days before immunization.(29) On days 5 and 10 after immunization, mice received another injection of the CD4 depletion antibody. Sera were collected from immunized mice on days 7 and 14 after immunization for analysis of antibody levels.

### Adoptive transfer

OVA-specific CD4^+^ T cells were purified from spleens of CD45.1 OT-II mice using the Naïve CD4^+^ T cell Isolation Kit (Miltenyi Biotec, 130-104-453) according to manufacturer’s instructions. Ten thousand purified CD4^+^ OT-II cells were adoptively transferred into C57BL/6J WT recipients by retro-orbital injection. Two days later, the recipient mice received a transfusion of 100 μl of stored HOD blood or 200 μl of the vaccine. Sera were collected at days 6 and 10 after transfusion or vaccination, respectively, for assessment of antibody levels.

### Antibody detection by ELISA

To eliminate day-to-day technical variability, all sera samples from a single experiment were batched, run and analyzed on the same day. Anti-HEL responses were measured by HEL-specific enzyme-linked immunosorbent assay (ELISA) as previously described.(31,33) Briefly, high binding polystyrene plates (Corning 9018) were coated overnight at 4°C with 10 ⎧g/ml HEL (Sigma-Aldrich) in PBS. Plates were then washed with wash buffer (0.05% Tween-20 in PBS) and incubated with blocking buffer (2% BSA and 0.05% Tween-20 in PBS). Serum samples were serially diluted (starting at 1:100) in blocking buffer and incubated in coated plates for 1 hour at room temperature. Horseradish peroxidase-conjugated goat anti-mouse IgG Fcγ specific antibody (Jackson ImmunoResearch) and anti-mouse IgM (Jackson ImmunoResearch) was then used as a secondary stain at a 1:5,000 dilution. For ELISA assays that employed a urea wash, 6.5 M urea in DI water was added for 10 minutes after the incubation with sera samples and before the secondary antibody was added. Plates were washed twice before and after incubation with urea. After the incubation with the secondary antibody, wells were developed using TMB substrate (KPL) and quenched with 2N H_2_SO_4_ after 10 min at room temperature. Optical densities were measured at 450 nm. End-point titers were calculated using GraphPad Prism through interpolation of the cutoff value from the fit of the optical density vs. (1/serum dilution) curve for each sample using the “plateau followed by one-phase exponential decay” model. The cutoff value was defined as the average plus three standard deviations of signals from background wells (i.e. signal values from wells incubated with blocking buffer alone).

### Germinal center B cell detection by flow cytometry

Spleens were harvested and dissociated using gentle mechanical disruption. Single cell suspension was subjected to RBC lysis with RBC lysis buffer (ThermoFisher), resuspended in growth medium (10% FBS in RPMI) and filtered through 40 µm cell strainers (Falcon). 2 million cells were used for staining. All staining antibodies were diluted in FACS buffer (2% FBS, 0.5% BSA, and 0.5% NaN_3_ in PBS). GC B cells were characterized and enumerated by flow cytometry. Splenocyte surface staining was performed for: Brilliant Violet 421^TM^ anti-mouse CD19 (BioLegend), Brilliant Violet 711^TM^ anti-mouse IgD (BioLegend), PE anti-mouse/human GL7 antigen (BioLegend), Brilliant Violet 605^TM^ anti-mouse CD4, Brilliant Violet 605^TM^ anti-mouse CD8 and Zombie Yellow Dye^TM^ (BioLegend). Surface staining was followed by fixing and permeabilization using the Transcription Factor Buffer Set (BD Pharmingen). Intracellular staining was performed with Alexa Fluor 647 conjugated mouse anti-BCL6 antibody (BD Pharmingen). Flow cytometry was performed using the Attune Nxt and data were analyzed using FlowJo and GraphPad Prism.

### Statistical analysis

Statistical analysis and graphing was performed with GraphPad Prism software. Mann-Whitney test was performed on groups of interest. Where appropriate, Kruskal-Wallis test was performed prior to comparing individual samples via Mann-Whitney tests. A value of P < 0.05 was considered to be statistically significant and assigned *, whereas P < 0.01, P < 0.001 and P < 0.0001 were assigned **, *** and ****, respectively.

For linear regression analysis of IgG titers vs. number of transferred OT-II cells data, both IgG titers and number of OT-II cells were logarithmically transformed and plotted in GraphPad Prism. The rate of change, or slope, in IgG titers in response to numbers of transferred CD4^+^ T cells was calculated using the “Simple Linear Regression” analysis in GraphPad Prism. A linear fit to logarithmically scaled data demonstrates a power law or exponential relationship between the two variables. The slope from a log-log plot linear fit represents the exponent in the power law relationship, rather than just a constant multiplier as from a fit to a linear plot.

## Results

### Transfused mice efficiently produce IgM, while class switching to IgG is limited

In our previous work utilizing the HOD mouse model, we showed that despite both vaccination and transfusion driving the generation of class-switched IgG antibodies, transfusion driven antibody responses were predominantly of lower-affinity and shorter duration.(4) Since the previous study did not include a direct comparison of the extent of class-switching between transfusion and vaccination, we first investigated the levels and kinetics of anti-RBC antibody production post-transfusion and directly compared it to antibody generation in response to vaccination (Fig 1A). WT mice were either transfused with stored HOD blood or vaccinated with a similar protein antigen (HEL-OVA) combined with Alum adjuvant, a commonly used component in vaccines. Sera samples were collected at various time points post-immunization and analyzed for antigen-specific antibody production using ELISA (Fig 1A). This approach allowed us to account for potential differences in antibody kinetics arising from the distinct routes of immunization - transfusion via intravenous (IV) injection and vaccination via intraperitoneal (IP) injection.

**Figure 1.**
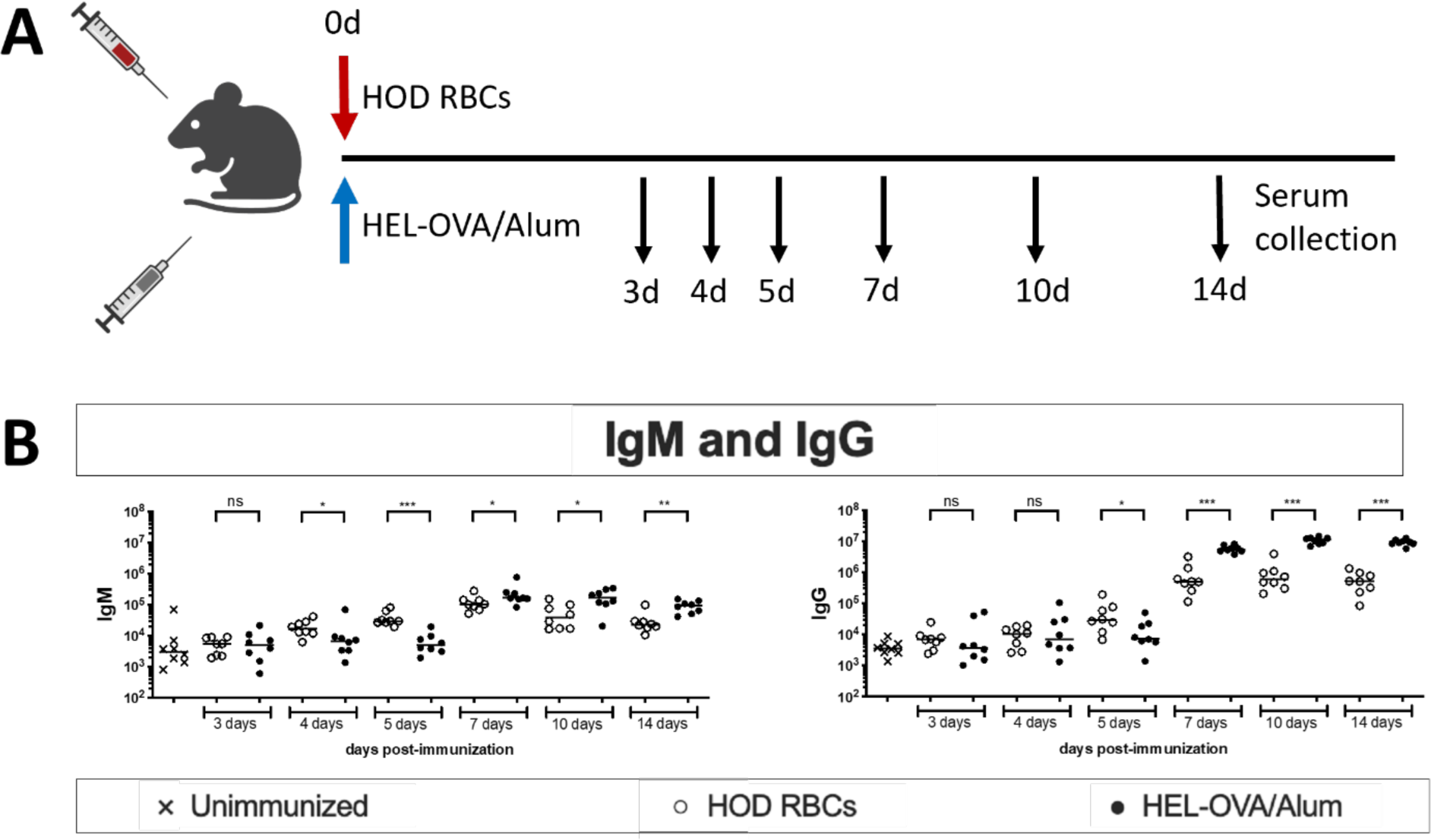
Transfused mice efficiently produce IgM, while class switching to IgG is limited. (A) Experimental setup. WT mice were either transfused with HOD RBCs or vaccinated with HEL-OVA/Alum. On days 3, 4, 5, 7, 10 and 14, sera were collected for assessment of antibody levels. (B) Anti-HEL IgM and IgG were measured by limiting dilution ELISA and presented as antibody titers. Each data point represents one mouse. Bars on scatter plots are median values. Figure shows a representative experiment out of 4. *P<0.05, **P<0.01, ***P<0.001, ****P<0.0001, ns P>0.5.

Transfused mice exhibited a more rapid response to immunization relative to vaccination, producing detectable levels of anti-HEL antibodies by 4-5 days post-transfusion (dpt), with peak antibody production observed at 7 dpt (Fig 1B). In contrast, vaccinated mice showed a relatively delayed response, with detectable anti-HEL antibodies appearing 1-2 days later compared to transfusion and reaching peak production between 10-14 days post-immunization (dpi). While anti-HEL IgM levels in both groups decreased shortly after reaching peak production, total IgG levels were more sustained. Specifically, IgG levels in both transfused and vaccinated mice remained stable from day 7 to day 14 post-immunization (Fig 1B).

Comparison of anti-HEL antibody levels between the two groups revealed that both transfused and vaccinated mice generated similar peak levels of anti-HEL IgM (Fig 1B). However, transfused mice produced higher levels of anti-HEL IgG at 5 dpi compared to vaccinated mice, reflecting differences in kinetics of antibody production between transfusion and vaccination. Surprisingly, despite this early IgG production, the peak levels of total IgG in transfused mice was significantly lower than that observed in vaccinated mice (Fig 1B). These findings suggest that B cells produce anti-HEL IgM with equal efficiency following transfusion and vaccination. However, in contrast to IgM production, class switching to IgG is more robust in vaccinated mice compared to transfused mice.

### IgG, but not IgM, production depends on CD4^+^ T cell help after transfusion

We next set out to determine the potential factors leading to less robust class switching to IgG in response to transfused RBCs relative to vaccination. Previous studies have consistently shown that strong antibody responses to various canonical immune stimuli, including protein-adjuvant vaccinations, depend on an intact CD4^+^ T cell compartment.(12,30) Given the well-established role of CD4^+^ T cells in facilitating the transition from IgM to IgG antibody production, we further investigated and directly compared their contribution to transfusion and vaccination induced IgG responses. To interrogate the role of CD4^+^ T cells in transfusion and vaccination driven antibody responses, we compared the effects of CD4^+^ T cell depletion prior to immunization on IgM and IgG production in transfused and vaccinated mice. Anti-HEL antibody levels were measured at 7 and 14 dpi to assess the impact of CD4^+^ T cell depletion (Fig 2A).

**Figure 2.**
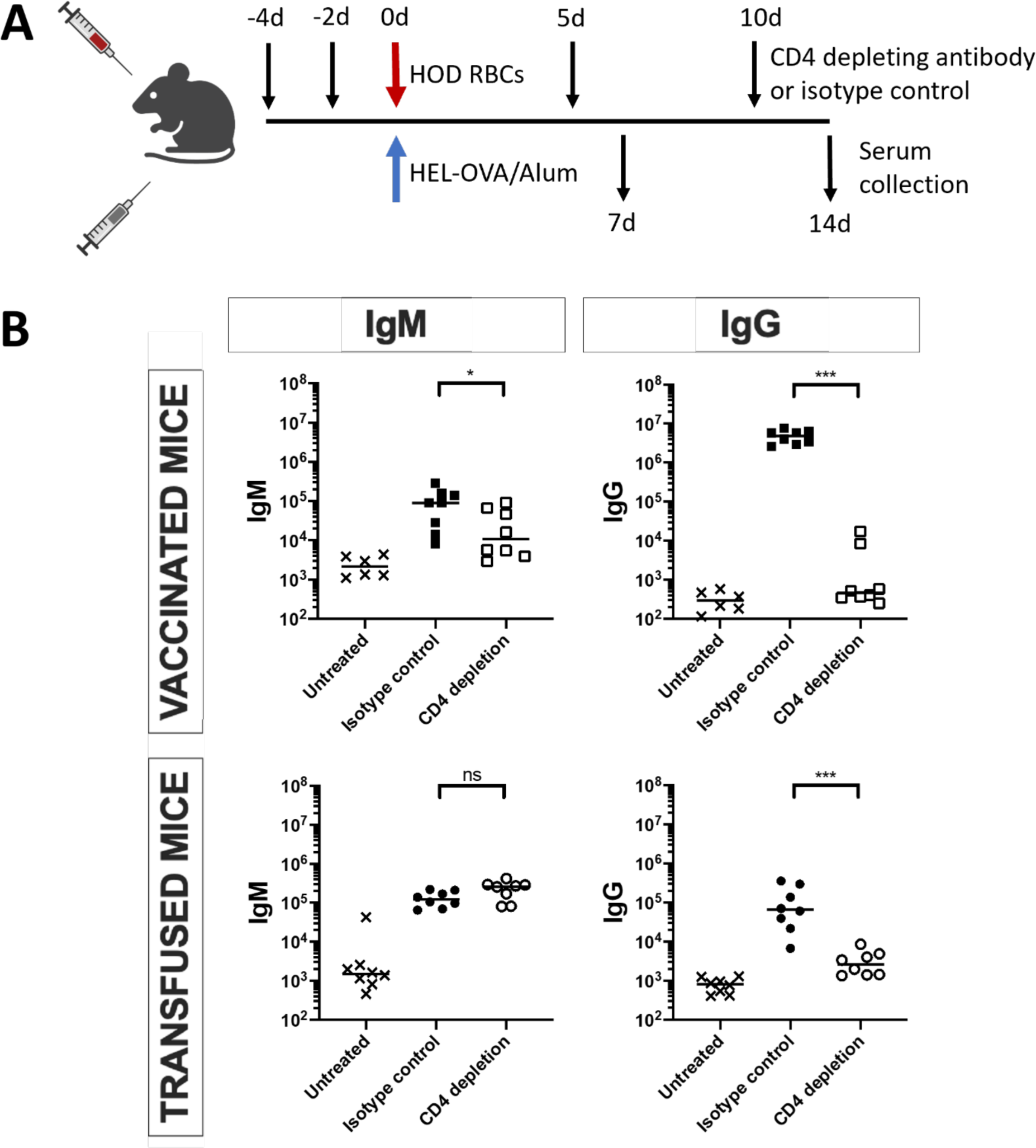
IgG, but not IgM, production depends on CD4^+^ T cell help after transfusion. (A) Experimental setup. WT mice received CD4^+^ depleting antibody 4 and 2 days before immunization and subsequently on days 5 and 10 after immunization. Sera were collected for assessment of antibody levels on days 7 and 14 after immunization. (B) Anti-HEL IgM and IgG were measured by limiting dilution ELISA and presented as antibody titers. Each data point represents one mouse. Bars on scatter plots are median values. Figure shows a representative experiment out of 3. *P<0.05, **P<0.01, ***P<0.001, ****P<0.0001, ns P>0.5.

Interestingly, we observed a differential impact of CD4^+^ T cell depletion on anti-HEL IgM production in response to transfusion and vaccination (Fig 2B). In vaccinated mice, CD4^+^ T cell depletion resulted in a significant reduction in IgM production, indicating that CD4^+^ T cells are essential for a robust IgM response post-vaccination. However, in transfused mice, CD4^+^ T cell depletion had no significant impact on IgM production, suggesting that IgM generation post-transfusion is independent of CD4^+^ T cell help. This indicates a fundamental difference in how IgM is produced in response to transfusion versus vaccination.

When examining IgG production, CD4^+^ T cell depletion led to a marked reduction in total IgG levels in both transfused and vaccinated mice (Fig 2B). This highlights the critical role of CD4^+^ T cells in facilitating class switching to IgG, regardless of immunization method. These results suggest that CD4^+^ T cell help is essential for effective class switching and IgG production in response to both transfusion and vaccination.

In summary, these findings show that while both IgM and IgG production after vaccination relies heavily on CD4^+^ T cell help, IgM production following transfusion is independent of CD4^+^ T cell help but CD4^+^ T cells still remain necessary for transfusion induced class switching to IgG.

### IgG, but not IgM, production depends on CD40-CD40L interaction after transfusion

Despite our results that both transfusion and vaccination driven IgG production critically requires CD4^+^ T cell help, the mechanisms that lead to relatively diminished IgG class switching in response to transfused RBCs remain unclear. We hypothesized that the lower levels of IgG class switching in response to transfusion with stored HOD RBCs are a result of suboptimal CD4^+^ T cell help provided to B cells post-transfusion compared to immunization with Alum and HEL-OVA vaccination. With the goal of better understanding and eventually manipulating the quality of CD4^+^ T cell help in response to transfusion with stored HOD RBCs, we next focused on investigating the molecular mechanisms that mediate CD4^+^ T cell help for B cell class switching to IgG.

CD40-CD40L signaling is a well-characterized and critical mechanism through which CD4^+^ T cells provide help to antibody producing B cells. The engagement of CD40 expressed on B cells with CD40L expressed on activated CD4^+^ T cells is known to be essential for IgG class switching and GC formation for T-dependent antigens in both mice and humans. We investigated the effects of blocking CD40L binding to CD40 on anti-HEL IgM and IgG production in response to stored HOD RBCs and compared it directly to the effects of CD40-CD40L blocking in response to HEL-OVA and Alum vaccination. We utilized a well-characterized monoclonal antibody (clone MR1) that antagonizes CD40-CD40L interactions *in vitro* and *in vivo*.(20,25,29,34) The CD40L blocking antibody was administered prior to immunization and subsequently twice per week up until 14 dpi. Sera samples were then analyzed for the production of anti-HEL antibodies (Fig 3A).

**Figure 3.**
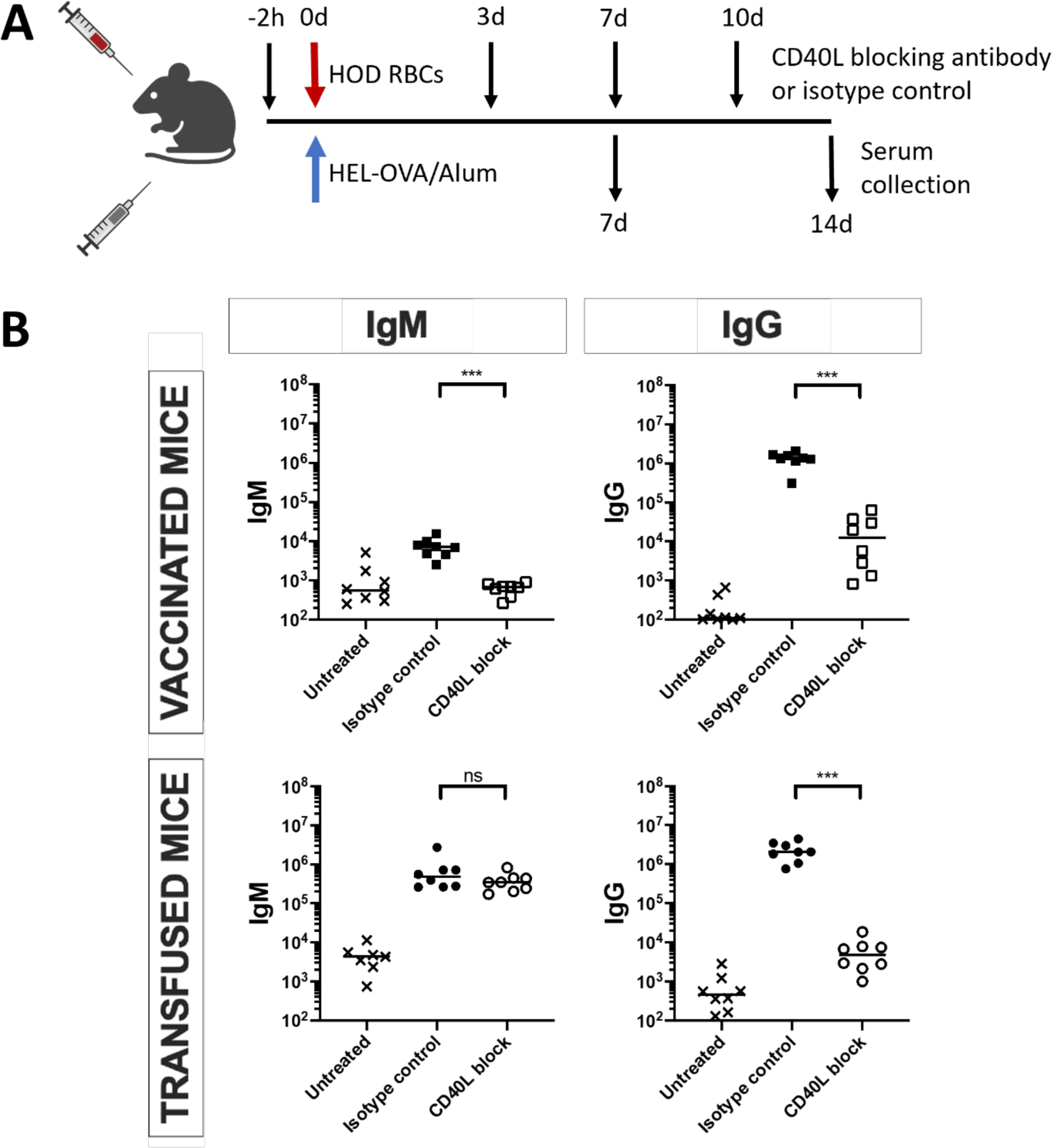
IgG, but not IgM, production depends on CD40-CD40L interaction after transfusion. (A) Experimental setup. WT mice received CD40L blocking antibody 2h before immunization and subsequently on days 3, 7 and 10 after immunization. Sera were collected for assessment of antibody levels on days 7 and 14 after immunization. (B) Anti-HEL IgM and IgG were measured by limiting dilution ELISA and presented as antibody titers. Each data point represents one mouse. Bars on scatter plots are median values. Figure shows a representative experiment out of 3. *P<0.05, **P<0.01, ***P<0.001, ****P<0.0001, ns P>0.5.

In vaccinated mice, application of the CD40L blocking antibody resulted in a significant reduction of anti-HEL IgM levels compared to mice receiving an isotype control (Fig 3B). The same treatment had no effect on IgM production in transfused mice, with anti-HEL IgM levels showing no significant difference between mice receiving the CD40L blocking antibody or an isotype control. In terms of IgG production, vaccinated mice exhibited reduced IgG levels when treated with the CD40L blocking antibody compared to the isotype control (Fig 3B). Similar to vaccinated mice, transfused mice showed a dramatic reduction in antigen-specific IgG levels following treatment with the CD40L blocking antibody.

Based on these results, we conclude that CD40-CD40L signaling is a necessary mechanism for CD4^+^ T cell help driven IgG class switching in response to both vaccination and transfusion. However, unlike vaccination, transfusion driven IgM responses were independent of CD40-CD40L interactions.

### Exogenous stimulation of CD40 enhances IgG production after transfusion

Given the necessity of CD40-CD40L signaling for IgG production in response to transfusion, we next investigated whether exogenous stimulation of the potentially suboptimal CD40-CD40L signaling induced by transfused stored HOD RBCs could be used to enhance IgG class switching. To stimulate CD40-CD40L signaling in B cells, we utilized an anti-CD40 agonistic antibody known to stimulate CD40 signaling in cells expressing CD40 on their surface.(35) We treated recipient mice with either the anti-CD40 agonistic antibody or an isotype control prior to transfusion with stored HOD RBCs or immunization with HEL-OVA and Alum vaccination, and compared the resulting anti-HEL antibody responses (Fig 4A).

**Figure 4.**
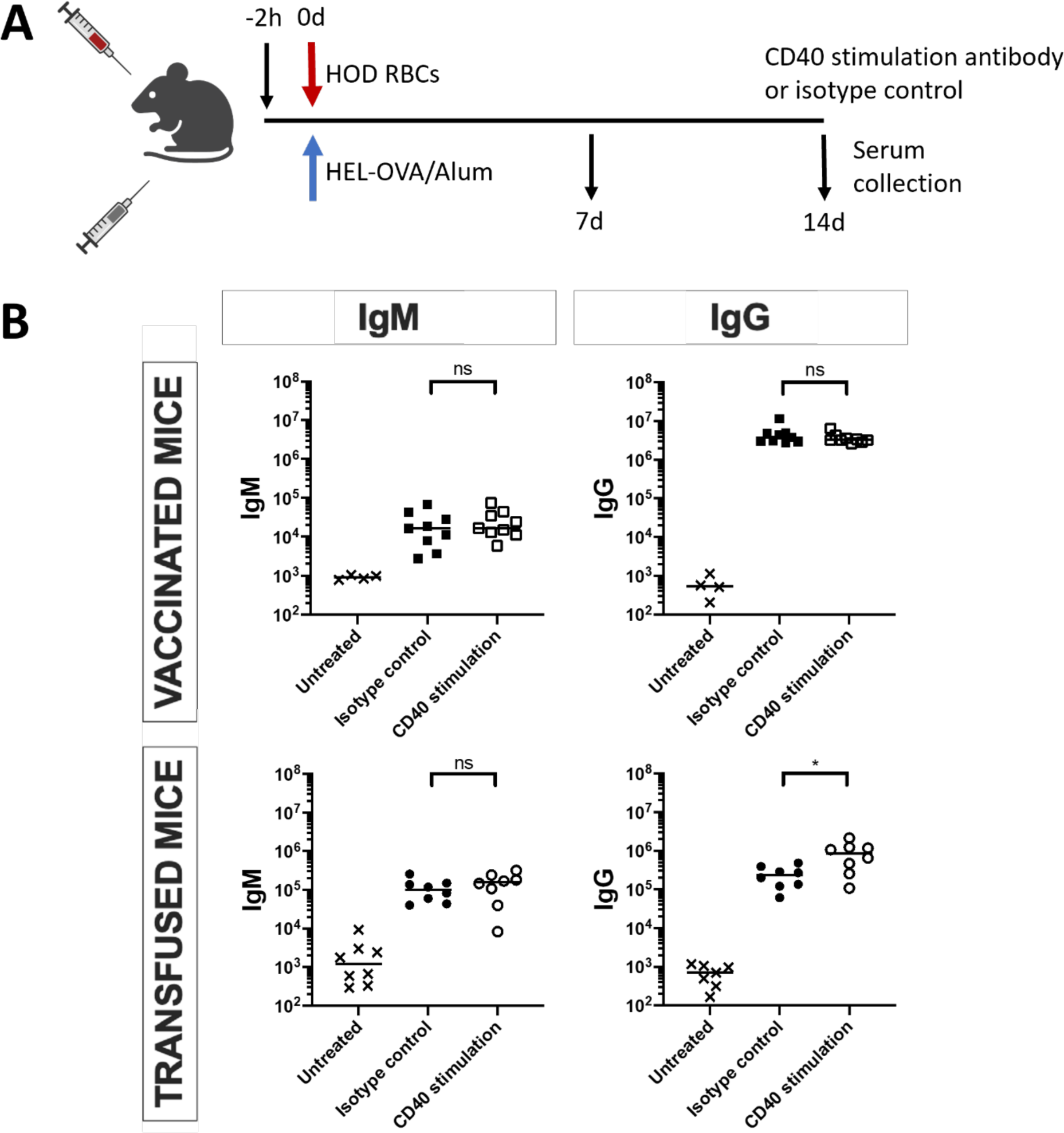
Exogenous stimulation of CD40 enhances IgG production after transfusion. (A) Experimental setup. WT mice received CD40 stimulation antibody 2h before immunization. Sera were collected for assessment of antibody levels on days 7 and 14 after immunization. (B) Anti-HEL IgM and IgG were measured by limiting dilution ELISA and presented as antibody titers. Each data point represents one mouse. Bars on scatter plots are median values. Figure shows a representative experiment out of 3. *P<0.05, **P<0.01, ***P<0.001, ****P<0.0001, ns P>0.5.

In vaccinated mice, CD40L-CD40 blocking reduced IgM production, but supplemental CD40 stimulation did not enhance IgM levels. IgM production remained comparable to that observed in mice treated with an isotype control antibody (Fig 4B). This suggests that additional CD40 signaling does not further boost IgM responses after vaccination, indicating that CD40-mediated help is not a limiting factor for IgM production in response to vaccination. In transfused mice, CD40 stimulation had no significant effect on IgM production, consistent with our loss-of-function findings (Fig 4B). This reinforces the observation that IgM generation after transfusion occurs independently of the CD40L-CD40 interaction.

In contrast, the impact of additional CD40 stimulation on IgG production revealed a stark difference in antibody responses between transfusion and vaccination (Fig 4B). In mice vaccinated with HEL-OVA and Alum, exogenous CD40 stimulation did not increase total IgG levels, indicating that CD40-mediated help during vaccination is already at saturating levels required for robust IgG production. Conversely, in mice transfused with stored HOD RBCs, CD40 stimulation significantly increased total IgG levels compared to mice that did not receive additional CD40 stimulation. These results confirm our hypothesis that CD40 signaling in response to transfusion, but not vaccination, is at a suboptimal level for class switching and IgG production.

Surprisingly, despite the significant increase in IgG production with exogenous CD40 stimulation in transfused mice, the total anti-HEL IgG levels were still approximately an order of magnitude lower compared to vaccinated mice (Figs 4B and 1B). This finding suggested to us that although CD40-CD40L signaling is necessary for the provision of CD4^+^ T cell help driven IgG production, it alone is not sufficient for driving maximal levels of class-switching to IgG in response to transfusion.

### Class switching to IgG in transfused mice depends on T cell help in a dose response manner

Since CD40 stimulation was not sufficient to drive IgG production in response to transfused HOD RBCs to the same levels as vaccinated mice, we next explored additional methods to enhance CD4^+^ T cell help in response to transfusion. In addition to CD40-CD40L signaling, CD4^+^ T cells help drive IgG production through multiple molecular mechanisms. As these mechanisms likely act in concert to drive CD4^+^ T cell dependent IgG production, we sought to directly enhance CD4^+^ T cell help in response to transfusion through adoptive transfer of additional antigen-specific T cells in recipient mice.

To achieve this goal, we utilized the OT-II mouse model, in which the majority of CD4^+^ T cells express a T Cell Receptor (TCR) specific to the OVA antigen.(33,36) As both HOD RBCs and the HEL-OVA protein used in our vaccine contain the OVA antigen, this model allowed us to precisely control the number of antigen-specific CD4^+^ T cells introduced into WT recipients and compare the resulting IgG production in response to transfusion and vaccination.

Lymphocyte precursor frequency has been shown to play an important role in shaping immune responses in mice.(37,38) We chose the range of transferred OT-II cell numbers in recipient mice to span from close to endogenous frequency of splenic OVA specific CD4^+^ T cells in C57BL/6 mice on the lower end to about two orders of magnitude higher numbers on the other end.(38,39) Prior to immunization, we purified OVA-specific CD4^+^ T cells from OT-II mice and transferred 1000, 10,000, or 100,000 OT-II cells into WT recipients, approximately corresponding to 2X, 20X, and 200X the endogenous frequency of OVA-specific CD4^+^ T cells respectively. After immunization with either stored HOD RBCs or HEL-OVA in Alum vaccination, sera were collected at the peak of antibody production, and anti-HEL antibody levels were measured and compared (Fig 5A).

**Figure 5.**
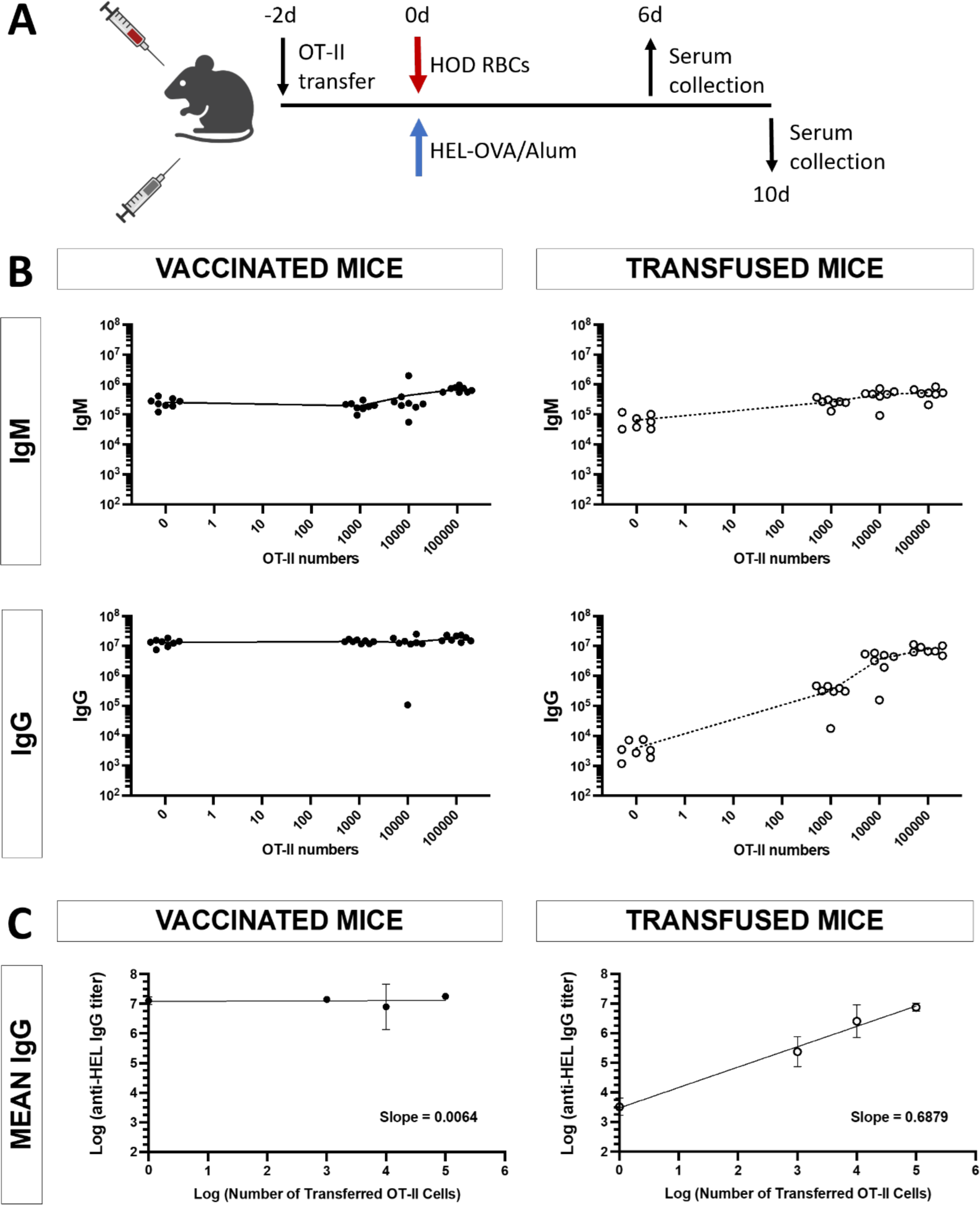
Class switching to IgG in transfused mice depends on T cell help in a dose response manner. (A) Experimental setup. WT mice received different numbers of OT-II cells 2 days before immunization. Sera were collected for assessment of antibody levels on days 10 and 6 after vaccination and transfusion, respectively. (B) Anti-HEL IgM and IgG were measured by limiting dilution ELISA and presented as antibody titers. Each data point represents one mouse. Bars on scatter plots are median values. Figure shows a representative experiment out of 2. *P<0.05, **P<0.01, ***P<0.001, ****P<0.0001, ns P>0.5. (C) Linear regression curve fit for anti-HEL IgG titers vs number of transferred OT-II cells on a log-log plot for vaccinated and transfused mice (same data as in panel B). Linear fit on a log-log plot represents a power law relationship between transferred OT-II cell numbers and IgG titers. Slope, which represents the rate of change in IgG titers in response to OT-II numbers, corresponds to the exponent in the power law equation.

In vaccinated mice, addition of antigen-specific CD4^+^ T cells did not enhance antibody production significantly (Fig 5A). IgM levels remained unchanged for up to the transfer of 10,000 OT-II cells, only showing a small increase in response to the transfer of 100,000 OT-II cells. Compared to mice that did not receive additional OT-II cells, no significant difference was observed in IgG titers even after the transfer of cell numbers that were 200X higher than the endogenous lymphocyte frequency. These results align with our earlier findings with CD40 stimulation, and confirm the hypothesis that CD4^+^ T cell help in response to vaccination is naturally at saturating levels.

In stark contrast, transfused mice exhibited a significant increase in antibody levels following the transfer of CD4^+^ T cells, even at the lowest dose of 1000 OT-II cells corresponding to ~2X the endogenous frequency (Fig 5B). Both IgM and IgG production increased in a dose-dependent manner, with the effect on IgG being more pronounced. Transferring 1,000 OT-II cells (~2X the endogenous frequency) led to less than a 5-fold increase in IgM, while IgG levels rose nearly 100-fold. At the highest dose of 100,000 cells (~200X the endogenous frequency), IgM levels increased less than 10-fold, while IgG levels increased by an astonishing 2,000-fold.

Linear regression analysis enabled us to quantitatively compare the dependence of IgG production to the number of transferred OT-II cells in transfusion versus vaccination (Fig 5C). We log-transformed both IgG titers and numbers of OT-II cells, then proceeded to fit a straight line to the log-log plot. A linear fit on a log-log plot is indicative of a power law relationship between the two variables, with the rate of change, or slope, representing the exponent value. Our fits to IgG vs. OT-II cell number plots generated slope values of ~0.69 for transfused mice and ~0.0064 for vaccinated mice (Fig 5C). The rate of change in IgG levels in response to transferred OT-II cells was two orders of magnitude higher in transfused mice compared to vaccinated mice. These analyses further support our findings that vaccination driven IgG production remains relatively unchanged in response to increased CD4+ T cell numbers, whereas transfusion driven IgG responses increase exponentially.

These findings suggest that transfusion does not inherently provide sufficient CD4^+^ T cell help for maximal antibody production, making CD4^+^ T cell help a limiting factor. Conversely, vaccination stimulates sufficient CD4^+^ T cell help, rendering additional assistance unnecessary.

### Supra-physiological numbers of CD4^+^ cells increase GC B cell differentiation and high-affinity IgG production in transfused mice

Class switching to IgG, particularly high-affinity IgG, is largely believed to result from B cells undergoing selection within GCs with the assistance of CD4^+^ T cells, although evidence suggests class switching can also occur outside of GCs. In our previous work using the HOD mouse model, we showed that transfusion with HOD RBCs fails to generate robust GCs and leads to significantly lower levels of high-affinity IgG production compared to vaccination.(4) Our gain-of-function experiments, using adoptive transfers of varying numbers of antigen-specific CD4^+^ T cells, revealed that increasing CD4^+^ T cell numbers significantly enhances class switching to IgG in a dose-dependent manner in transfused mice, while this effect was not observed in vaccinated mice. Given these results, we determined to further investigate whether the adoptive transfer of antigen-specific CD4^+^ T cells also affected GC formation and generation of high-affinity IgG antibodies in response to transfusion.

We examined GC differentiation and the production of high-affinity IgG in transfused versus vaccinated mice receiving different numbers of OT-II cells (Fig 6A). In transfused mice, increasing the number of antigen-specific CD4^+^ T cells led to a corresponding increase in GC differentiation and high-affinity IgG production, as measured by urea-based ELISAs to denature low-affinity IgGs. In contrast, even the addition of supra-physiological numbers of CD4^+^ T cells had no significant impact on GC differentiation or high-affinity IgG production in vaccinated mice (Fig 6B).

**Figure 6.**
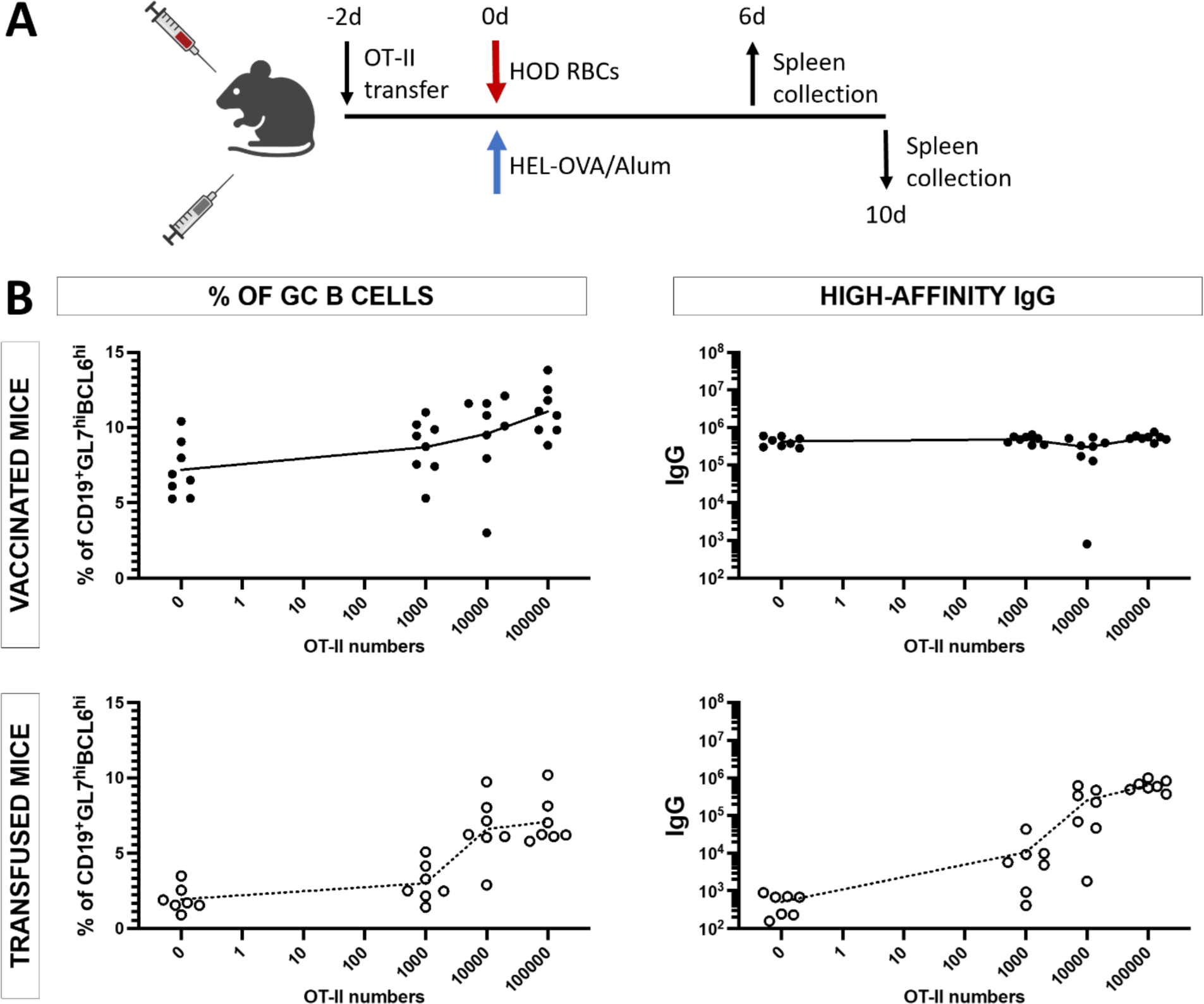
Supra-physiological numbers of CD4^+^ cells increase GC B cell differentiation and high-affinity IgG production in transfused mice. (A) WT mice received different numbers of OT-II cells 2 days before immunization. Spleens and sera were collected for assessment of GC B cell differentiation and high-affinity IgG on days 10 and 6 after vaccination and transfusion, respectively. (B) GC B cells were quantified using flow cytometry. Anti-HEL IgG were measured by limiting dilution ELISA, that incorporated urea incubation, and presented as antibody titers. Each data point represents one mouse. Figure shows a representative experiment out of 2. *P<0.05, **P<0.01, ***P<0.001, ****P<0.0001, ns P>0.5.

These findings highlight that CD4^+^ T cell help is the limiting factor for efficient class switching, development of robust GC reactions, and high-affinity IgG production in the context of transfusion-induced immune responses.

## Discussion

Development of class-switched anti-RBC IgG alloantibodies remains a significant clinical problem for chronically transfused patient populations. Despite recent advances, immune responses to antigens on transfused RBCs remain incompletely characterized. In our previous work, we showed that unlike a canonical immune stimulus such as protein-adjuvant vaccination, transfusion fails to generate robust GCs and results in predominantly short-lived and low-affinity IgG antibodies.(4) In the current study, we further characterized immune responses against transfused RBCs through direct comparison of class-switching and CD4^+^ T cell help between transfusion with stored HOD RBCs and vaccination with HEL-OVA emulsified in Alum.

While both transfusion and vaccination led to equivalent IgM production, IgG antibody levels generated in response to transfusion were significantly lower compared to vaccination. To our knowledge, these data are the first direct comparison of antibody responses against the same antigen presented via either transfused RBCs or adjuvant-based vaccination. Our results indicate that in addition to preferential production of short-lived and low-affinity IgG antibodies, the overall extent of class-switching to IgG is significantly reduced in response to transfusion compared to vaccination. Investigation into potential mechanisms that underlie the relatively diminished IgG class-switching in response to transfusion revealed clear differences between the role of CD4^+^ T cells in transfusion and vaccination driven antibody responses. CD4^+^ T cell depletion led to a significant reduction in vaccination driven IgG and IgM production, whereas only IgG production was significantly inhibited in transfused mice with IgM levels remaining unaffected. Blocking CD40-CD40L signaling in recipient mice recapitulated the effects of CD4^+^ T cell depletion equivalently, demonstrating the necessity of CD40-CD40L interactions in mediating CD4^+^ T cell help to B cells. These findings reveal a distinct reliance on CD4^+^ T cell help in the overall mechanisms underlying antibody production in response to transfusion versus vaccination. The role of CD40-CD40L signaling has previously been reported as important in antibody responses to transfused RBCs in mice.(29) However, in that model the mice were transfused with fresh HOD RBCs along with poly(I:C), a potent TLR3 agonist that leads to robust activation of inflammation and innate immune responses. Here we show that CD40-CD40L interactions are necessary for IgG production in response to stored HOD RBCs as well. Though the CD4^+^ T cell dependence of IgG production in response to stored HOD RBCs has been reported previously (30), a direct comparison CD4^+^ T cell help in regulating IgM and IgG production in response to transfusion and vaccination clearly illustrates the non-canonical nature of transfused RBCs as an immune stimulus.

Since both vaccination and transfusion driven IgG production depend critically on CD4^+^ T cell help, we hypothesized that differences in the ability to stimulate CD4^+^ T cells may be the limiting factor underlying relatively diminished class-switching to IgG in transfused mice. As CD4^+^ T cell help is also known to be essential for robust GC reactions and high-affinity IgG production, CD4^+^ T cell help is likely a limiting factor in the comparatively lower development of these responses in transfused mice as well. Effects of providing additional CD4^+^ T cell help on antibody production in transfused mice supports this hypothesis. Exogenous stimulation of CD40 signaling resulted in a significant increase in transfusion driven anti-HEL IgG production but failed to reach equivalent titers to vaccinated mice. Adoptively transferring varying numbers of OVA-specific CD4^+^ T cells prior to transfusion with HOD RBCs resulted in a significant dose-dependent increase in anti-HEL IgG antibodies, approaching equivalent titers to vaccination driven IgGs. Concomitant with a significant increase in class-switching to IgG, transfused mice also showed a dose-dependent increase in GC development and high-affinity IgG production in response to additional CD4^+^ T cell transfer also approaching equivalent levels to vaccinated mice. In stark contrast, vaccination driven total IgG production, GC development, and high-affinity IgG titers remained unaffected in response to CD40 stimulation and adoptive transfer of OVA-specific CD4^+^ T cells. Immune responses in vaccinated mice remained unchanged even after provision of CD4^+^ T cell numbers corresponding to about two orders of magnitude increase in endogenous frequency of OVA specific CD4^+^ T cells, suggesting that protein-adjuvant vaccination is able to stimulate saturating levels of CD4^+^ T cell help at physiological CD4^+^ T cell numbers. Whereas in transfused mice, merely doubling the OVA-specific CD4^+^ T cell frequency through adoptive transfer led to a significant increase in antibody responses and GC development. Collectively, our findings using the HOD mouse model suggest that the comparatively reduced degree of class-switching to IgG, robust GC reactions, and high-affinity IgG production in response to transfusion are a result of sub-optimal stimulation of CD4^+^ T cell help compared to vaccination and can be “rescued” by exogenous provision of additional CD4^+^ T cell help.

Despite being necessary for CD4^+^ T cell help driven IgG production in response to transfusion, CD40 stimulation alone was not sufficient to increase anti-HEL IgG titers equivalent to that generated in response to vaccination. Efficient CD4^+^ T cell help to B cells is critically mediated through multiple cell-cell contact dependent interactions via receptor-ligand pairs such as, but not limited to, CD40-CD40L, ICOS-ICOSL, and SAP-SLAM.(11,15,16,19,26,34) Along with relevant intracellular signaling triggered by these cell-cell contact molecules themselves, they also allow for selective directional secretion of key effector cytokines produced by CD4^+^ T cells that stimulate antibody production from B cells.(17,18) It is likely that while blocking CD40-CD40L interaction leads to disruption of physical interactions between CD4^+^ T cells and B cells, thus preventing provision of CD4^+^ T cell help, enhancing CD40 stimulation alone is unable to replicate the contribution of multiple other critical mediators of CD4^+^ T cell help. Though the enhancement of CD4^+^ T cell help may require simultaneous stimulation of multiple effector pathways, our results with CD40-CD40L blocking suggest that singular disruption of other cell-cell contact mediators between T and B cells may be effective at significantly inhibiting anti-RBC IgG production. Going forward, investigating the contribution of other critical cell-cell contact dependent mediators of CD4^+^ T cell help can generate multiple pharmaceutical targets for prevention of RBC alloimmunization in chronically transfused patients.

Anti-HEL IgM production in response to transfusion with stored HOD RBCs was unaffected by both CD40-CD40L blocking and CD4^+^ T cell depletion, whereas IgG production was significantly inhibited. This intriguing feature of antibody responses against transfused RBCs may be explained by differences in specific B cell populations regulating antibody production between transfusion and vaccination. Previous work has shown that antibody responses to transfused stored HOD RBCs are primarily generated by marginal zone B cells (MZBs).(30) Unlike follicular B cells, MZBs are localized in the splenic marginal zone and display features of both innate and adaptive immunity.(40,41) The anatomical positioning of MZBs allows them to effectively capture and react to blood-borne antigens, leading to rapid antibody generation that provides critical protection during the time it takes for the follicular B cells to produce long-lived high-affinity antibody responses.(40–42) The predominant role of MZBs in anti-HOD antibody generation could also explain the relatively rapid antibody production in response to transfusion compared to vaccination observed in this study. In response to canonical T-independent antigens, MZBs can produce both IgM and IgG antibodies without CD4^+^ T cell help.(40,41) However, in response to transfused RBCs only IgM production was independent of CD4^+^ T cell help while being critically required for IgG production. Thus, antibody production in response to transfused RBCs display features of both canonical T cell-dependent and T cell-independent (TI) immune responses, preventing their classification as exclusively TD or TI antigens. Collectively, these results further demonstrate the non-canonical nature of transfused RBCs as an immunological stimulus.

Provision of additional CD4^+^ T cell help via adoptive transfer of OT-II cells to recipient mice enhanced GC reactions and high-affinity IgG production in response to transfusion. The mechanisms through which additional numbers of CD4^+^ T cells enhance these features remain unclear. Unlike antibody responses to vaccination which rely critically on CD4^+^ T cell help to follicular B cells (5,7), transfusion driven antibody responses have been reported to be regulated exclusively by MZB cells.(30) In response to canonical T-dependent antigens, MZB cells receive help from CD4^+^ T cells to generate GC reactions and high-affinity IgG antibodies.(40,43,44) It is also possible that exogenously increasing the number antigen-specific CD4^+^ T cells allows these cells to now interact with follicular B cells to trigger the increased GC reactions and high-affinity IgG production. Whether the enhancement of GCs and high-affinity IgG production in response to increased CD4^+^ T cell numbers in transfused mice is a result of increased T cell help to MZB cells, increased engagement of follicular B cells, or a combination of both remains to be investigated. Individually depleting MZBs or follicular B cells along with provision of additional CD4^+^ T cells will greatly help illuminate the mechanisms underlying effects of increased CD4^+^ T cell frequency on antibody responses in transfused mice.

We have shown that the lack of robust GC reactions and diminished high-affinity IgG generation in response to transfusion are a result of sub-optimal CD4^+^ T cell help stimulated by transfused RBCs. Another feature of transfusion induced antibody responses, reported in our previous work, is that they are predominantly short-lived relative to vaccination.(4) Whether the inability of transfused RBCs to generate long-lived antibody responses is also a result of sub-optimal CD4^+^ T cell help remains to be tested. RBC antibody evanescence can have significant clinical consequences in chronically transfused patient populations.(45–47) Going forward, it would be interesting to measure the duration of antibody responses in transfused mice after the adoptive transfer of increasing doses of CD4^+^ T cell help.

The question remains as to why CD4^+^ T cell help is stimulated sub-optimally in response to HOD RBC transfusion versus vaccination. Antigen presenting cells, specifically dendritic cells, play a critical role in the activation of CD4^+^ T cells via presentation of antigenic peptides via MHC-II receptors.(48,49) Due to differences in the route of immunization, antigen persistence, and the biochemical characteristics of immune stimuli between transfusion with stored HOD RBCs and HEL-OVA/Alum, the populations and functional states of dendritic cells may also vary in response to transfusion and vaccination. Multiple studies have shown that antibody responses to transfused RBCs critically require the spleen.(50–52) In future work, we plan to investigate splenic dendritic cell populations that interact with transfused stored RBCs and assess their functional properties and the nature of CD4^+^ T cell activation.

Patient alloimmunization responses to transfused RBCs are known to be highly antigen-dependent, with a great range of immunogenicity observed clinically for different blood group antigens.(1) This clinically relevant aspect of RBC alloimmunization is also evident in mouse models, where recipient mice exhibit differences in both the extent of alloimmunization and the underlying cellular and molecular mechanisms across different model RBC antigens. (1,53) Given this caveat, the results of the current study are likely relevant to understanding alloimmunization responses to a subset of clinically important RBC antigens but may not represent immune responses to all blood group antigens. As a next step, it will be essential to perform similar experiments using mouse models engineered with different RBC antigens.

In summary, our results expand on our previous work in characterizing the immune responses against transfused HOD RBCs. We show that compared to protein-adjuvant vaccination, transfused HOD RBCs stimulate distinct antibody responses in terms of kinetics, extent of class-switching, GC reaction development, and high-affinity IgG production. These differences between transfusion and vaccination can be explained by their engagement of different B cell populations and differences in the degree of CD4^+^ T cell help stimulation. Going forward, we plan on investigating the role of CD4^+^ T cell help in anti-RBC alloantibody evanescence and mechanisms underlying sub-optimal CD4^+^ T cell help stimulated by transfused RBCs. Results from the current study extends ongoing efforts to better understand non-canonical immune responses to transfused RBCs, and are essential for developing therapeutic interventions to treat and prevent the clinical consequences of RBC alloimmunization in chronically transfused patients.

